# Particle size distribution and optimal capture of aqueous macrobial eDNA

**DOI:** 10.1101/001941

**Authors:** Cameron R. Turner, Matthew A. Barnes, Charles C.Y. Xu, Stuart E. Jones, Christopher L. Jerde, David M. Lodge

## Abstract

1. Detecting aquatic macroorganisms with environmental DNA (eDNA) is a new survey method with broad applicability. However, the origin, state, and fate of aqueous macrobial eDNA - which collectively determine how well eDNA can serve as a proxy for directly observing organisms and how eDNA should be captured, purified, and assayed - are poorly understood.
2. The size of aquatic particles provides clues about their origin, state, and fate. We used sequential filtration size fractionation to measure, for the first time, the particle size distribution (PSD) of macrobial eDNA, specifically Common Carp (hereafter referred to as Carp) eDNA. We compared it to the PSDs of total eDNA (from all organisms) and suspended particle matter (SPM). We quantified Carp mitochondrial eDNA using a custom qPCR assay, total eDNA with fluorometry, and SPM with gravimetric analysis.
3. In a lake and a pond, we found Carp eDNA in particles from >180 to <0.2 μm, but it was most abundant from 1-10 μm. Total eDNA was most abundant below 0.2 μm and SPM was most abundant above 100 μm. SPM was ≤0.1% total eDNA, and total eDNA was ≤0.0004% Carp eDNA. 0.2 μm filtration maximized Carp eDNA capture (85%±6%) while minimizing total (i.e., non-target) eDNA capture (48%±3%), but filter clogging limited this pore size to a volume <250 mL. To mitigate this limitation we estimated a continuous PSD model for Carp eDNA and derived an equation for calculating isoclines of pore size and water volume that yield equivalent amounts of Carp eDNA.
4. Our results suggest that aqueous macrobial eDNA predominantly exists inside mitochondria or cells, and that settling plays an important role in its fate. For optimal eDNA capture, we recommend 0.2 μm filtration or a combination of larger pore size and water volume that exceeds the 0.2 μm isocline. *In situ* filtration of large volumes could maximize detection probability when surveying large habitats for rare organisms. Our method for eDNA particle size analysis enables future research to compare the PSDs of eDNA from other organisms and environments, and to easily apply them for ecological monitoring.

## INTRODUCTION

Environmental DNA (eDNA) is DNA extracted from bulk environmental samples (e.g., soil, water, air) without isolating target organisms or their parts from the sample. The concept and the term both originate from microbiology (Ogram *et al.* 1987) where the target DNA in environmental samples is from abundant live and dead microbes. In contrast, *macrobial* eDNA is the DNA of large organisms such as animals or plants that occurs in environmental samples. Although macrobial eDNA has been studied since 1991 in fields such as human forensics (Hochmeister *et al.* 1991), agricultural transgenics (Widmer *et al.* 1997), paleogenetics (Willerslev *et al.* 2003), and fecal pollution source tracking (Martellini *et al.* 2005), it was only in 2008 that eDNA was first used to detect aquatic macrofauna (Ficetola *et al.* 2008). Aqueous macrobial eDNA has garnered particular interest as a simple and sensitive alternative to directly observing rare aquatic macrofauna. Despite a recent burst of papers demonstrating its use for detection of diverse macrofauna, we have much to learn about the origin (e.g., primary physiological source), state (e.g., intra- or extra-cellular), and fate (e.g., suspension time) of aqueous macrobial eDNA (Lodge *et al.* 2012). These three domains collectively determine how well eDNA can serve as a proxy for directly observing organisms, and how eDNA should be captured to make robust inferences about organism presence or abundance. In this paper we determine the size distribution of particles containing eDNA and draw size-based inferences about the state and fate of eDNA.

Martellini *et al.* (2005) first demonstrated that natural waters contain mitochondrial DNA (mtDNA) from vertebrates inhabiting the watershed. Their research focused on terrestrial mammals for which aqueous eDNA was presumably of allochthonous fecal origin. Autochthonous aqueous eDNA from aquatic macrofauna was subsequently described for amphibians (Ficetola *et al.* 2008), fish (Jerde *et al.* 2011), mammals, crustaceans, insects (Thomsen *et al.* 2012a), birds (Thomsen *et al.* 2012b), and reptiles (Piaggio *et al.* 2013) in lentic, lotic, and marine waters. eDNA-based surveys for rare fish or amphibians are more sensitive than traditional methods (Jerde *et al.* 2011; Thomsen *et al.* 2012b; Dejean *et al.* 2012) and eDNA disappears from surface water within 1 to 25 days of organism absence (Dejean *et al.* 2011; Thomsen *et al.* 2012a; b). Aqueous macrobial eDNA concentration is very low (up to 33 mtDNA copies mL^−1^) and correlates with organism density, at least for amphibians in small ponds and streams (Thomsen *et al.* 2012a; Pilliod *et al.* 2013) and fish in experimental ponds (Takahara *et al.* 2012). Aqueous macrobial eDNA has been captured (i.e., concentrated from water) by precipitation/centrifugation (Martellini *et al.* 2005; Ficetola *et al.* 2008), chromatography (Douville *et al.* 2007), lyophilization (Poté *et al.* 2009), ultrafiltration (pore size <0.1 μm; Takahara *et al.* 2012), and microfiltration (pore size >0.1 μm; Martellini *et al.* 2005; Goldberg *et al.* 2011; Jerde *et al.* 2011; Minamoto *et al.* 2011).

Despite the considerable variation in methods used to capture aqueous macrobial eDNA (Table S1), we are unaware of any systematic comparison between methods or description of the captured particles. The size of aquatic particles helps determine their characteristics and interactions with organisms, other particles, and the environment (Burd & Jackson 2009). Thus the size of aquatic particles containing macrobial eDNA is foundational to our understanding of the origin, state, and fate of this material. For example, large particles sink faster than small particles (Maggi 2013), so eDNA-based surveys aimed at determining very recent and local organism presence might need to target the upper end of the eDNA-containing particle size distribution (PSD). The PSD of aqueous macrobial eDNA can also inform decisions about sampling effort (e.g., water volume) and particle capture size (e.g., filter pore size). Currently these decisions are made using only precedent, trial and error, and logistical constraints. Finally, the PSD of macrobial eDNA may help mitigate one of the most troublesome issues associated with genetic testing for low copy number DNA in environmental samples: sample interference (Schrader *et al.* 2012). Non-target material suspended in water, including non-target DNA (Thompson *et al.* 2006), can interfere with eDNA recovery (Lloyd *et al.* 2009) and genetic assays (Hedman & Rådström 2013). If the PSD of macrobial eDNA differs from that of non-target material then size-based, selective enrichment of target eDNA may be possible.

Here we produce the first measurement of PSD for aqueous macrobial eDNA. We sequentially filtered water from a lake and pond inhabited by Common Carp (*Cyprinus carpio* Linnaeus, 1758, hereafter Carp) to create a series of size fractions containing the particles retained at each size, from ≥180 μm to <0.2 μm. In aquatic microbiology, 0.2 μm is the filter pore size below which aqueous eDNA exists as extracellular molecules free in solution (Matsui *et al.* 2004; Maruyama *et al.* 2008), and the vertebrate mitochondrion measures 0.2-8 μm (Flindt 2006 p. 254). Thus our size fractions spanned from extracellular and extraorganellar DNA (hereafter extramembranous; <0.2 μm) to particles larger than most vertebrate cells (≥180 μm). For each size fraction, we measured the concentration of Carp eDNA, total eDNA (DNA of any type), and suspended particle matter (SPM). This allowed us to compare PSDs among these three nested types of aquatic matter and evaluate the opportunity for size-based target enrichment.

## MATERIALS & METHODS

### Target species

We selected Carp as the target macrobial species because fish species, including Carp, have been the target of many previous studies of aqueous macrobial eDNA (Dejean *et al.* 2011; Jerde *et al.* 2011; Minamoto *et al.* 2011; Takahara *et al.* 2012, 2013; Thomsen *et al.* 2012a; b; Collins *et al.* 2012; Wilcox *et al.* 2013). In addition, Carp is listed at number 30 on the IUCN list of the world’s worst invasive species (Lowe *et al.* 2004). Because Carp inhabit many continents, our method for particle size analysis can be applied globally to assess the consistency of one species’ eDNA PSD across environmental conditions.

### Species-specific qPCR assay for eDNA quantification

We designed a species-specific SYBR Green qPCR assay to measure the concentration of only Carp eDNA in environmental samples. The qPCR primers were manually designed in an alignment of all available mitochondrial control region (D-loop) sequences from NCBI’s GenBank for Carp and the most closely related species that potentially co-occur in North America: Goldfish (*Carassius auratus*), Grass Carp (*Ctenopharyngodon idella*), Black Carp (*Mylopharyngodon piceus*), Crucian Carp (*Carassius carassius*), and Bigheaded Carp (*Hypophthalmichthys* spp.). We conducted further *in silico* testing of species specificity using NCBI Primer-BLAST tool. A set of candidate primers was selected for *in vitro* testing on tissue-derived total genomic DNA from Carp, Goldfish, Grass Carp, Black Carp, and Bigheaded Carp. We selected one primer set (Ccrp_Dlp_F: GAGTGCAGGCTCAAATGTTAAA; Ccrp_Dlp_R: GTAAGGATAAGTTGAACTAGAGACAG; 146 bp amplicon) that demonstrated efficient amplification from Carp and no amplification from any of the non-target species. Finally, we conducted *in situ* testing of this assay using eDNA samples from natural water bodies with and without Carp. The assay did not amplify eDNA from a small lake without Carp (Knop Lake, Indiana, USA). The assay amplified eDNA from two large lakes with Carp (Lakes Michigan and Huron, Michigan, USA), and Sanger sequences (ABI 3730xl, Applied Biosystems) of these qPCR products matched the target amplicon from Carp. The DNA melting temperature (T_m_) of all qPCR products from all eDNA samples matched that from tissue-derived Carp DNA, as determined by a melting curve.

During development and application of the assay, we used 20 μL reactions containing 10 μL of Power SYBR Green Master Mix (Life Technologies), 4 μL of DNA, 2 μL of 4 μg*μL^−1^ non-acetylated BSA (Life Technologies), and 2 μL of each 3 μM primer. We performed all reactions on an Eppendorf Mastercycler ep realplex2 S thermocycler with the following reaction conditions: 95°C for 10 min.; 55 2-step cycles of 95°C for 15 sec. and 60°C for 1 min., 95°C for 15 sec., 60°C for 15 sec., and a 20 min. melting curve up to 95°C for 15 sec. Fluorescence data collection occurred during the 1 min. 60°C step and during the melting curve.

We used absolute quantification qPCR with a standard curve to quantify Carp eDNA. We made the standard curve DNA by amplifying the entire mtDNA control region (1022 bp) (Liu & Chen 2003) from tissue-derived Carp DNA. After confirming a single gel electrophoresis band approximately 1000 bp in size we purified this PCR amplicon using ExoSAP and quantified it using 5 μL of PCR product with a Qubit fluorometer (Invitrogen) and the Qubit dsDNA High Sensitivity kit. We converted from DNA weight to DNA copies using the median molecular weight of the 95% consensus 1022 bp amplicon sequence from all *C. carpio* mitogenomes on GenBank (631214 g*mole^−1^) as calculated by OligoCalc (Kibbe 2007). We stored single use 7.7E + 06 copies*μL^−1^ aliquots of this standard DNA at −20°C. qPCR standard curves were prepared by serial 10-fold dilution from 7.7E + 06 copies*μL^−1^ down to 1 copy*μL^−1^ in low TE buffer (10 mM Tris, 0.1 mM EDTA) and stored at 4°C while in use (Bellete *et al.* 2003). All manipulations of standard DNA, eDNA, and reagents for qPCR used low bind tubes and low bind aerosol barrier pipette tips (Ellison *et al.* 2006).

### Water bodies

We collected water from two lentic water bodies, selected to provide a strong contrast in simplicity of the physical environment, in macrofaunal diversity, and in abundance of the target species (Carp). At Potawatomi Zoo in South Bend, Indiana, USA (41.670629, −86.216634), we sampled a small (0.06 ha, maximum depth 2 m) concrete-lined outdoor pond with flow-through treated municipal water that contained approximately 500 large adult Carp. We expected that this extremely high Carp density would provide sufficiently high carp eDNA concentration to allow quantification in every size fraction following sequential filtration. The more natural, lower Carp density environment was St. Mary’s Lake at the University of Notre Dame, Indiana, USA (41.701497, −86.244312), a natural spring-fed mesotrophic lake (10.7 ha, maximum depth of 9.1 m). We lack a quantitative estimate of Carp density for this lake but based on our many visual observations, the density of Carp is far lower in the lake than in the zoo pond.

### Water collection and sampling

In December 2011 (“winter”) we conducted our primary sampling event for each water body. In April 2012 (“spring”) we conducted a supplemental sampling event at St. Mary’s Lake because Carp eDNA concentrations measured in winter were very low (see Results and Discussion). For each sampling event we pumped 12 L of water from ≤1 m depth into an autoclaved carboy using autoclaved silicone peristaltic tubing with internal diameter of 9.5 mm (Masterflex L/S 36) and a portable, battery-powered peristaltic pump (Alexis, Pegasus Pump Company, Bradenton, Florida, USA). At the pond we pumped water from the east end of the pond where Carp were aggregated by extending the tubing approximately 2 m from shore. At the lake we pumped water from a small embayment on the south end as we rowed a bleach-decontaminated boat to and from a point approximately 100 m from shore. Carboys were transported to the laboratory within approximately 20 minutes and water was distributed into autoclaved 300 mL bottles. We stored the 300 mL bottles on ice until each was used for particle size fractionation, which began immediately and finished within 10 hours.

### Particle size fractionation

Filtration can bias particle size analysis by retaining particles smaller than the filter pore size or passing particles larger than the pore size (Droppo 2006). However, our analysis required chemical assay of the SPM to quantify its subcomponents, total eDNA and Carp eDNA. Thus particle sizing methods that do not require fractionation (e.g., electrical, optical, or flow cytometry particle counting) were not applicable (Reynolds *et al.* 2010). Filtration was also the most appropriate method given our aim to use PSDs for informing optimal eDNA capture – which is typically done by filtration. For these reasons we used sequential filtration size fractionation (Figure 1) to generate the PSDs for SPM, total eDNA, and Carp eDNA. We followed the recommendations of Droppo (2006) to minimize filtration artifacts: low vacuum pressure, small water volume, and filters with precise pores. We used nylon net filters (180, 100, and 60 μm mesh size; Millipore) and track-etched polycarbonate (PCTE) filters (20, 10, 1, 0.2 μm pore size; GE Osmonics) because these provide the sharpest separation between particle sizes and the best agreement between nominal and effective pore size (Sheldon 1972; Droppo 2006). By contrast, ‘tortuous path’ type filters such as glass-fiber and cellulose membrane actually have a wide array of effective pore sizes in every filter, rather than well-defined pores (see Droppo 2006 for a detailed discussion). Thus our PSDs, while accurate, may not translate directly to the nominal pore sizes of tortuous path filters, and we recommend the use of PCTE filters for future particle size analysis of aqueous macrobial eDNA.

**Figure 1.**
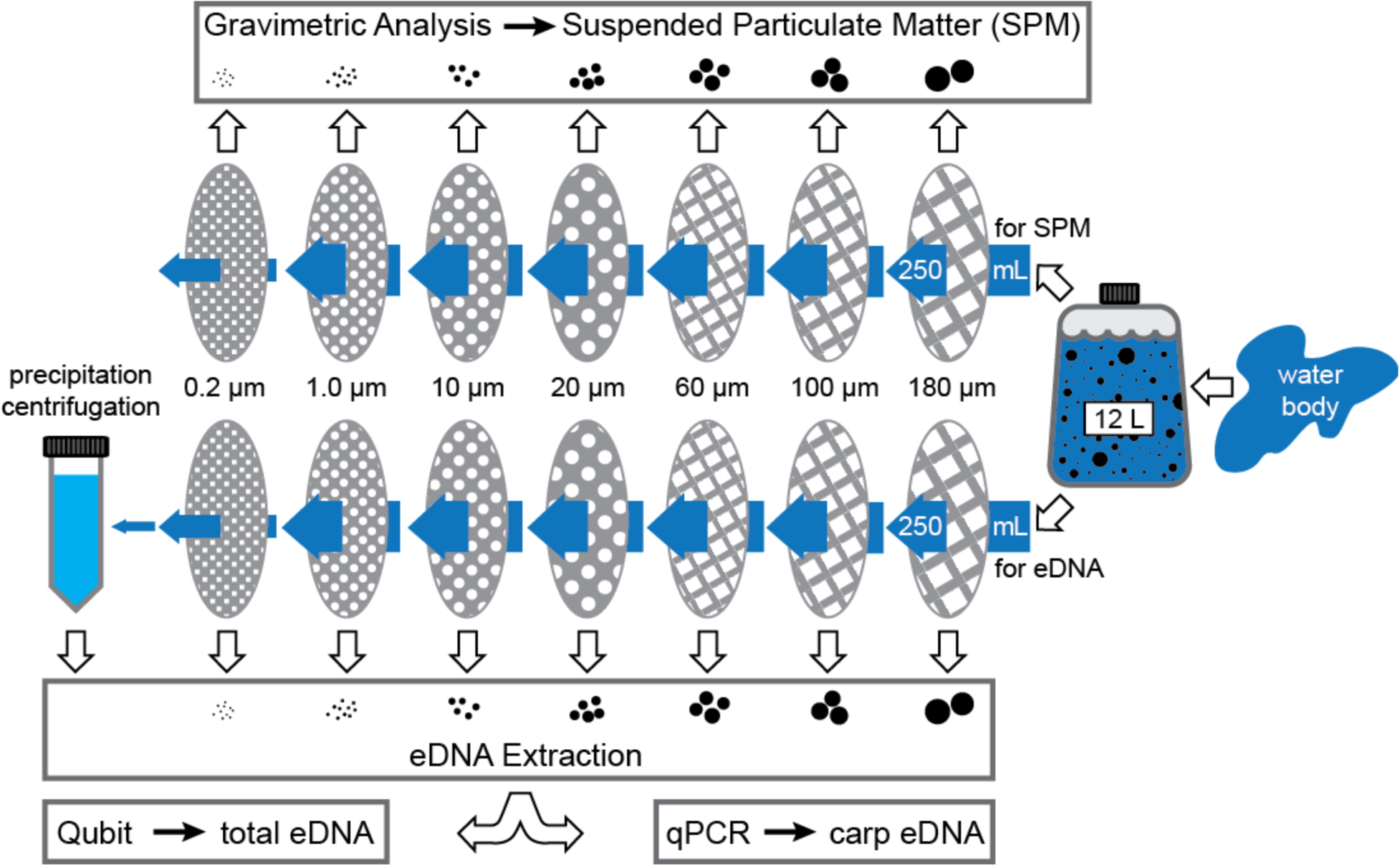
Diagram of particle size fractionation by sequential filtration. Surface water from throughout the water body was pumped into a 12 L carboy. 250 mL samples from this carboy were size fractionated by sequential filtration through filters of decreasing pore size, 180–0.2 μm. For each water body ten 250 mL samples provided filter retentate used in gravimetric analysis while ten samples were used for eDNA extraction and eDNA assays. Samples used for eDNA also provided a 15 mL subsample of 0.2 μm filtrate from which eDNA was precipitated and pelleted by centrifugation. Only 150 mL of 1.0 μm filtrate was used for filtration through the 0.2 μm filter to avoid filter clogging.

From each of the ten (winter) or three (spring) bottles, 250 mL was sequentially filtered through 47 mm diameter filters of decreasing pore size using gentle vacuum pressure (<64 cm Hg). We used 47 mm diameter magnetic filter funnels (Pall) and 1 L polypropylene vacuum flasks (Nalgene) that were decontaminated between uses by immersion in a 10% bleach solution followed by copious rinsing with reverse osmosis treated water. To avoid inaccurate size fractionation caused by filter clogging (which we observed in a pilot experiment), we used only 150 mL of the 1 μm filtrate for filtration through the 0.2 μm filter. 15 mL of the 0.2 μm filtrate was transferred to a 50 mL centrifuge tube containing 33.5 mL of 100% ethanol and 1.5 mL of 3 M sodium acetate then frozen at −20°C for subsequent eDNA capture following the precipitation procedure of Ficetola *et al.* (2008) with a centrifuge force of 3220 g. We placed used filters in CTAB buffer (Coyne *et al.* 2005) and stored them at −20°C until subsequent DNA extraction.

For the primary sampling event (winter) only, we conducted size fractionation for gravimetric analysis at the same time and in the same manner as for DNA analysis except all filters were individually pre-desiccated, pre-weighed and stored in individually labeled petri dishes. Following filtration, a filter disc was carefully returned to its petri dish retentate side up and placed in a 60°C desiccation chamber for 24 hours prior to gravimetric analysis. The final size fraction (<0.2 μm) was not included in gravimetric analysis because we were unable to weigh the tiny pellet produced by precipitation/centrifugation of the 0.2 μm filtrate.

### Assaying eDNA: contamination precautions

We conducted eDNA extraction, total eDNA quantification, and qPCR setup in a strictly pre-PCR laboratory located on a separate floor and opposite end of the building from our post-PCR laboratory. eDNA extraction and total eDNA quantification were performed on a bench used exclusively for those purposes and qPCR setup was performed on a different bench used exclusively for that purpose which is located on the opposite end of the pre-PCR laboratory (Mifflin 2007). A DNA extraction negative control was included for every filter pore size and for every batch of extractions. Each qPCR plate included three no template control (NTC) reactions.

### Assaying eDNA: eDNA extraction

We extracted eDNA from filters and precipitation pellets following the CTAB extraction of Coyne *et al.* (2005). PCTE filters dissolve during the extraction and nylon net filters were shredded in their extraction tubes using flame- and bleach-sterilized scissors. Final eDNA pellets were re-suspended in 100 μL of low TE buffer.

### Assaying eDNA: total eDNA quantification (Qubit)

We quantified total eDNA using 5 μL of eDNA extract in the Qubit dsDNA High Sensitivity kit and a Qubit fluorometer (Invitrogen). The limit of detection for this method is 0.5 ng*mL^−1^ in the 200 μL assay tube (corresponding to 0.02 ng*μL^−1^ in our eDNA extracts) thus we set the eDNA concentration at 0.02 ng*μL^−1^ for eDNA extracts with concentrations too low for Qubit measurement. This reflects our assumption that eDNA existed at every size fraction, even if the sample volume limitations of our size fractionation produced some extract concentrations too low for Qubit detection.

### Assaying eDNA: Carp eDNA quantification (qPCR)

We measured Carp eDNA concentration in each eDNA extract using triplicate reactions of the qPCR assay performed as described above. To minimize variation between replicate reactions caused by imperfect pipetting of small eDNA extract volumes, we combined eDNA extract and master mix (all other reagents) for three reactions into one tube then dispensed to three plate wells using an electronic repeating pipette (Xplorer 5-100 μL, Eppendorf). Each qPCR plate included a five point standard curve from 3.1E + 04 copies*reaction^−1^ down to 3 copies*reaction^−1^. The fluorescence threshold for each plate was automatically determined by the Eppendorf realplex software using the default Noiseband setting. The fluorescence baseline was calculated for every reaction individually using the default Automatic Baseline setting of the Eppendorf realplex software. Every amplification profile and melting curve profile was visually examined to confirm exponential amplification and a T_m_ matching that of standard curve reactions. Following the recommendation of Ellison *et al.* (2006) for low copy number qPCR, we calculated concentrations for each reaction, assigning zero concentration to non-detect reactions and averaging concentration across the three technical replicates for an eDNA extract. In some reactions the measured reaction copy number was slightly below one so we rounded all reaction copy numbers up to the next largest integer.

### Assaying SPM (gravimetric analysis)

Desiccated filters containing captured particles were weighed using a microbalance. To eliminate the effect of static charge buildup on microbalance readings, we passed every filter through two 2U500 polonium-210 ionizers held in a Staticmaster positioner (Amstat Industries, Inc., Glenview, Illinois, USA).

### Data conversion, analysis, and modeling

To compare the PSD shape of Carp eDNA, total eDNA, and SPM, we converted concentration in each size fraction to percent of total (Figure 2). This assumed that the final fractionation step (precipitation for Carp eDNA and eDNA, 0.2 μm filtration for SPM) captured all remaining matter for each of the three types, respectively. To evaluate the optimal capture of Carp eDNA, we converted concentration of Carp eDNA, total eDNA, and SPM to cumulative percent captured (capture efficiency, E) at each fractionation step. To allow the most general evaluation possible from our experiment, we pooled the lake and pond capture efficiency data (winter only) without making any assumptions about the similarity or difference between the two water bodies. To estimate Carp eDNA capture efficiency as a continuous function of filter pore size (i.e., a continuous PSD for Carp eDNA) we modeled our discrete capture efficiency data with the complementary cumulative distribution function (CCDF) of the Weibull distribution, a flexible distribution that was originally developed and is widely used to describe PSDs (Rosin & Rammler 1933; Zobeck *et al.* 1999; see Appendix S1 for details). Finally, we used this continuous model to calculate ‘isoclines’ of filter pore size and filtered water volume where the Carp eDNA PSD estimates identical captured amounts of Carp eDNA.

**Figure 2.**
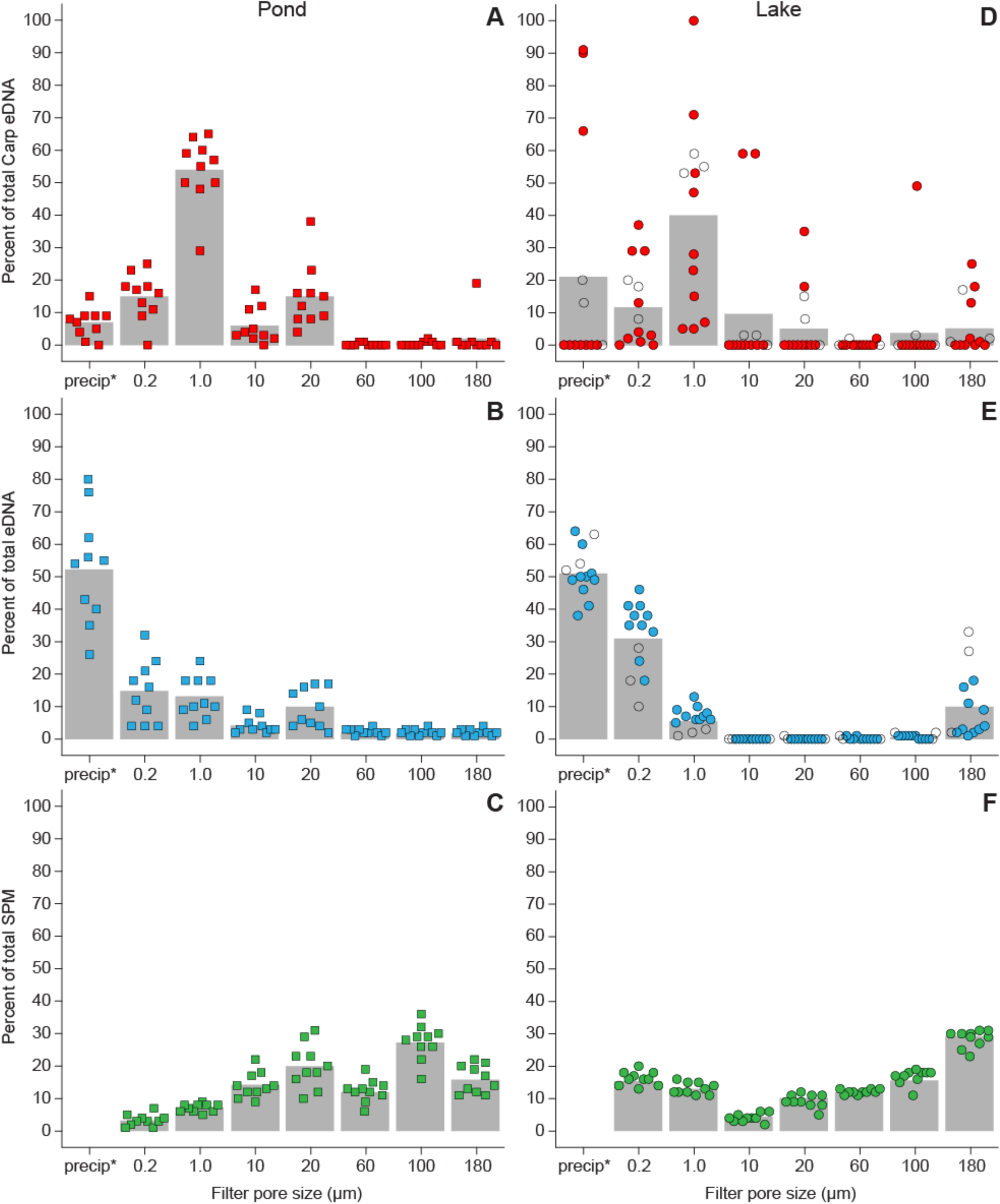
Percent of total carp eDNA, total eDNA, and SPM in each particle size fraction per water sample from the pond (A, B, C, respectively) and lake (D, E, F, respectively). *The final fractionation step was DNA precipitation using the 0.2 μm filtrate. Filled symbols are winter; open symbols are spring (only the lake was sampled in spring; SPM was not measured in spring). Data points are horizontally ‘jittered’ within each size fraction to reduce visual overlap of data points. Grey bars show the average value for each size fraction, which includes the supplemental spring data from the lake in (D) and (E).

To calculate the proportion of total eDNA that was made up by Carp mtDNA, we converted the qPCR-measured Carp eDNA copy number to weight of Carp mitogenomes. This was done using the molecular weight of one arbitrarily chosen Carp mitogenome from NCBI GenBank (10244181 g*mole^−1^; accession number: JN105352.1) as calculated by OligoCalc (Kibbe 2007).

## RESULTS

All extraction negative controls and qPCR NTCs tested negative for Carp eDNA. In both water bodies, particles containing Carp eDNA were detected in all of the size fractions, from ≥ 180 μm to < 0.2 μm, and Carp eDNA was most abundant in the 1-10 μm particle size fraction (Figure 2). On average there was more Carp eDNA in particles larger than 0.2 μm for both water bodies (pond: 10 of 10 replicates; lake: 7 of 10 in winter, 3 of 3 in spring) although this difference was significant only for the pond (Chi-square goodness of fit test; pond: χ^2^(1, n = 10) = 10.00, p < .01; lake-winter:χ^2^(1, n = 10) = 1.60, p = 0.21; lake-all: χ^2^(1, n = 13) = 3.77, p = 0.05). The average percent of Carp eDNA in particles larger than 0.2 μm was 93%±1% (mean ± s.e.m; pond), 75%±13% (lake-winter), and 78%±10% (lake-all). These results demonstrate that Carp eDNA can exist in very large particles and suggest that extramembranous Carp eDNA (particles <0.2 μm) represents a small portion of the total size distribution of particles containing Carp eDNA. In contrast, total eDNA was most abundant in the <0.2 μm size fraction, and SPM was most abundant in the 100-180 μm size fraction (pond) or the ≥ 180 μm size fraction (lake) (Figure 2).

The average percent of SPM that was total eDNA was 0.01%±0.001% (mean ± s.e.m.; pond) and 0.1%±0.01% (lake) when summed across all size fractions. The average percent of total eDNA that was Carp mitochondrial eDNA was 0.0004%±0.0001% (pond) and 0.0000004%±0.0000001% (lake) when summed across all size fractions. These results demonstrate that total eDNA and SPM were nearly 100% non-target material with respect to the target, Carp mtDNA.

Filtration with a 0.2 μm pore size maximized Carp eDNA capture efficiency (85%±6%; cumulative data pooled across lake and pond) while minimizing total eDNA capture efficiency (48%±3%). Unfortunately, 0.2 μm pore size filters clog with very small throughput volumes (e.g., we were unable to filter 250 mL of 1.0 μm filtrate through our 0.2 μm filter). Maximum throughput volume can be increased by either increasing filter surface area or increasing filter pore size. Our data cannot provide guidance for increasing filter surface area, but we used the continuous PSD of Carp eDNA to estimate the combinations of pore size and volume that yield identical amounts of Carp eDNA. The estimated scale and shape parameters (λ and k) from the Weibull CCDF model (i.e., the continuous PSD) of Carp eDNA were 11.85 and 0.351 and the estimated Carp eDNA capture efficiency at 0.2 μm (E_0.2_) was 78.8% (Figure S1).

Thus the equation for estimating isoclines relative to 0.2 μm filtration is derived as follows: C* V_0.2_ *E_0.2_ = C*V_x_ *E_x_ (eqn 1), where C is the environmental concentration of the target, V is throughput volume, and E is target capture efficiency. Solving for V_x_ yields V_x_ = (V_0.2_*E_0.2_) / E_x_ (eqn 2). V_0.2_ is whatever throughput volume the researcher determines is feasible for a 0.2 μm pore size filter, E_0.2_ is target capture efficiency at 0.2 μm, and E_x_ is the continuous PSD of the target, yielding the final equation:

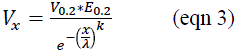

Figure 3 plots isoclines using the continuous PSD for Carp eDNA and several hypothetical values of V_0.2_.

**Figure 3.**
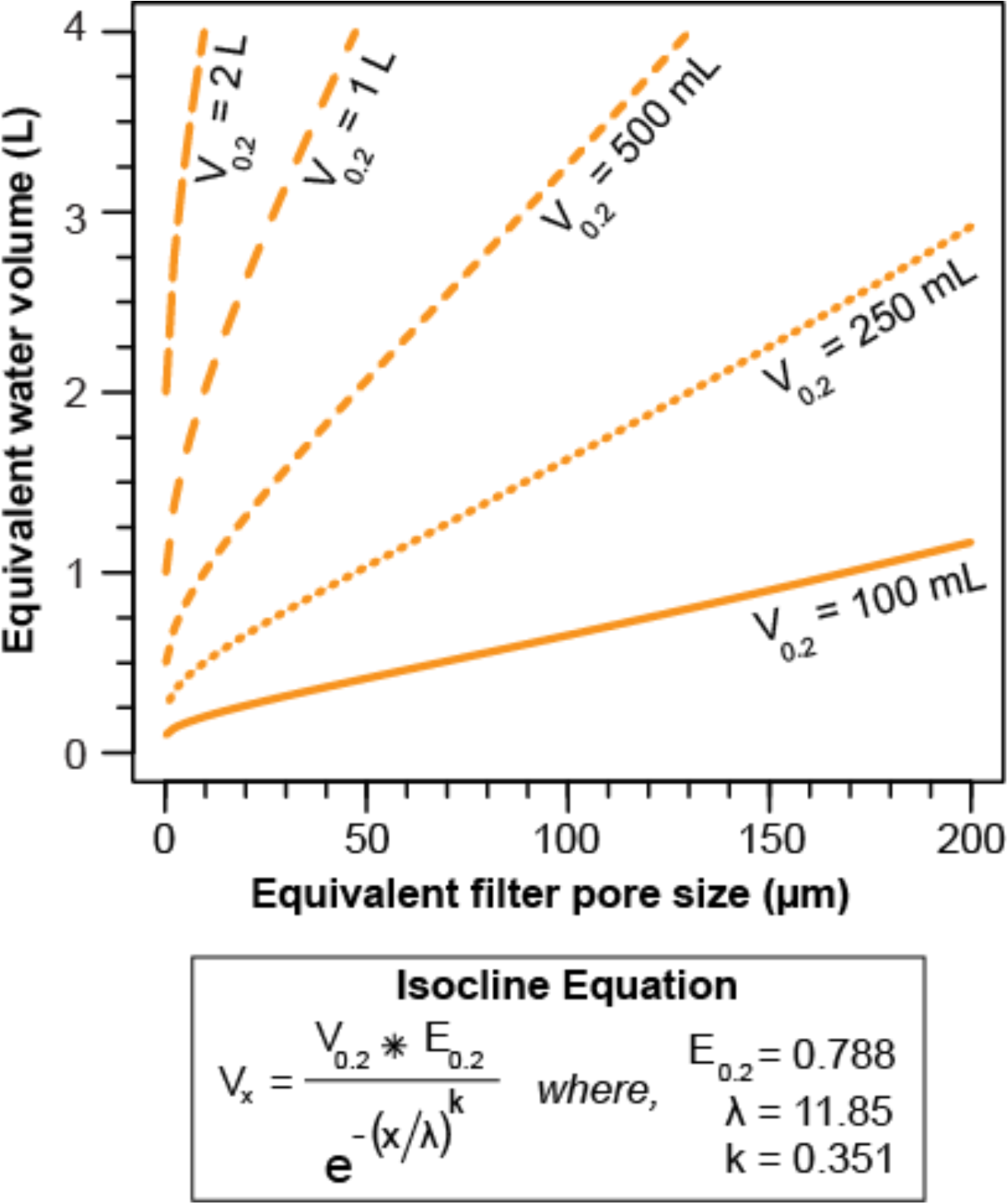
Isoclines showing combinations of filter pore size and water volume where the particle size distribution (PSD) of Carp eDNA predicts identical amounts of Carp eDNA captured. Isoclines are shown for five hypothetical examples of maximum throughput water volume for a 0.2 μm filter pore size (V_0.2_). The equation for calculating isoclines is shown using the 0.2 μm capture efficiency (E_0.2_) we estimated for Carp eDNA by fitting a Weibull CCDF to our cumulative size fractionation data; the scale (λ) and shape (k) parameters from the Weibull CCDF model of Carp eDNA PSD are also used in this equation (see Results and Appendix S1 for details).

## DISCUSSION

### Methodological considerations

As this paper represents the first investigation of particle size for aqueous macrobial eDNA, we first consider some methodological points that will be important should other scientists use our approach. The per reaction copy numbers of Carp eDNA measured by qPCR were very low, with a heavily right-skewed distribution and a mode of zero for both water bodies (Figure S2). This result is consistent with detailed empirical studies of qPCR applied to low copy number DNA (i.e., < 100 target copies per reaction) (Ellison *et al.* 2006). Research using qPCR to quantify eDNA should report how ‘non-detects’ (negative reactions from technical replicate sets where at least one reaction was positive) are handled. As demonstrated by Ellison et al. (2006), these non-detect reactions should be assigned a zero target concentration and included when averaging across technical replicates, as this provides better estimates of unknown sample concentration.

Even with this low copy number protocol applied, the frequency of negative reactions (79%) and non-detects (22%) in the lake-winter sampling event was high. The large sampling error (Figure 2) caused by these extremely low target DNA levels led us to conduct the supplemental spring sampling at the lake, because we expected higher fish activity in the spring would increase Carp eDNA concentration. Indeed, the frequency of negative reactions and non-detects dropped to 44% and 19%, respectively, in the lake-spring sampling event. The spring PSD from the lake supported our conclusions that Carp eDNA exists predominantly in particles larger than 0.2 μm and is most abundant between 1 and 10 μm (Figure 2). The supplemental spring sampling event also confirmed that total Carp eDNA concentration in the lake (summed across all size fractions) had increased four-fold since winter (from 1726 ± 602 to 7144 ± 1165 copies*L^−1^). These data represent the first description of seasonal change in eDNA concentration for aquatic macrofauna and underline the importance of reporting eDNA sampling dates, especially when comparing eDNA sampling with organism sampling (Thomsen *et al.* 2012a).

### State, and fate of aqueous macrobial eDNA

The majority of Carp eDNA was found in particles larger than 0.2 μm in both the pond (93%) and the lake (75%-78%), and Carp eDNA was most abundant in the 1-10 μm particle size fraction (Figure 2). In contrast, total eDNA was most abundant in the <0.2 μm size fraction and SPM was most abundant in the 100-180 μm (pond) or the <180 μm size fraction (lake) (Figure 2). Future research from other seasons, environments, and species will determine the generality of these results, but even at this early stage it seems appropriate to consider how the PSD can shed light on the state and fate of aqueous macrobial eDNA.

Aqueous eDNA from living macrofauna most likely originates as urine and feces, epidermal tissues and secretions, and reproductive cells and fluids. Much of this source material (e.g., feces) will enter the water column as large particles (>1000 μm), yet we generally found little Carp eDNA in particles larger than 60 μm. Therefore we conclude that these sources of eDNA must rapidly settle or break apart. Indeed, particle size is a major determinant of settling velocity, even for complex biological and biomineral aggregates (Maggi 2013). It is also well-documented that animal feces contain viable epithelial cells (10^−1^−10^6^ cells*g^−1^) and large amounts of the animal’s mtDNA (10^−1^−10^7^ copies*g^−1^) (reviewed in Caldwell *et al.* 2011) and that feces from aquatic macrofauna rapidly sink (Robison *et al.* 1981; Wotton & Malmqvist 2001). Thus at least one obvious source of aquatic macrobial eDNA appears to spend very little time suspended in the water column. We measured PSDs in still water whereas particle settling should be slower in flowing water, such as streams.

On the small end of the size spectrum there was generally little extramembranous Carp eDNA (<0.2 μm), at least as indicated by the 146 bp mtDNA region our qPCR assay targeted. One might predict an accumulation of smaller particles as discharged materials decompose, ultimately leading to an abundance of free eDNA. However, free eDNA may be exceptionally susceptible to hydrolysis given its full exposure to microbial extracellular enzymes that are abundant in aquatic systems (Matsui *et al.* 2001). Thus the abundance of Carp eDNA in the 1-10 μm fraction likely reflects a state of eDNA persisting within small cells and/or mitochondria. We cannot exclude the possibility that Carp eDNA in particles larger than 0.2 μm is actually extramembranous DNA aggregated or adsorbed onto larger particles (Burd & Jackson 2009; Suzuki *et al.* 2009), but the fact that animal mitochondria range in diameter from 0.2-1.2 μm and in length from 1-8 μm (Flindt 2006) suggest mitochondria as a parsimonious explanation for the pattern we observed. Indeed, during the regular apoptotic shedding of epithelial cells, intact mitochondria are released within apoptotic bodies and mtDNA is protected from the endonuclease degradation that rapidly degrades nuclear DNA (Murgia *et al.* 1992; Tepper & Studzinski 1993). For aquatic animals, this process will release whole mitochondria into the water column where their double membrane may resist lysis while the mitochondrial nucleoid further protects mtDNA (Rickwood & Chambers 1981).

The PSD of Carp eDNA indicates that settling, in addition to degradation, plays an important role in the fate of aqueous macrobial eDNA. Particles larger than 1 μm settle in natural waters (Isao *et al.* 1990) and 71%±5% of particles containing Carp eDNA exceeded this size (Figure 2). SPM in the 1-100 μm size range settles at terminal velocities between 0.4 mm*hr^−1^ and 40 m*hr^−1^, depending on size, shape, and composition (Maggi 2013). Thus it appears that most aqueous macrobial eDNA is in a constant state of downward flux. Most research on the fate of aqueous macrobial eDNA has not considered settling (Martellini *et al.* 2005; Kortbaoui *et al.* 2009; Dejean *et al.* 2011; Thomsen *et al.* 2012a), except one experiment that applied water circulation to eliminate it (Thomsen *et al.* 2012b). Our findings emphasize that degradation of DNA molecules and particle settling combine to reduce aqueous eDNA concentration over time and future research should try to measure both processes. Past eDNA degradation experiments may have underestimated the persistence of aqueous eDNA given that settled particles can return to the water column through bioturbation and water turbulence (Bloesch 1995).

We did detect a non-trivial amount of Carp eDNA in particles smaller than 1 μm (Figure 2). This fraction of aqueous macrobial eDNA is likely to remain in suspension indefinitely until the DNA molecules degrade (Isao *et al.* 1990). Water could advect eDNA in this state over long distances, especially in lotic systems, strong currents, and when free eDNA is protected via adsorption to other submicron particles (Saeki *et al.* 2011). For example, transgenes from genetically modified corn were detected in river water up to 82 km downstream of a corn cultivation plot (Douville *et al.* 2007), and plant DNA fragments as long as 1000 bp are detectable in groundwater (Poté *et al.* 2009). The stability and advection of free DNA in aquatic environments is sufficient that scientists use synthetic DNA as a hydrologic tracer (Sabir *et al.* 2002; Foppen *et al.* 2011). Future work examining the transport and persistence of naturally occurring aqueous macrobial eDNA across long distances will improve our understanding of the spatiotemporal window for inferring organism presence. The size of this window may depend on the particle size fraction that is tested for eDNA.

### Optimal capture of aqueous macrobial eDNA

Filtration with a 0.2 μm pore size was the best strategy for maximizing Carp eDNA and minimizing non-target eDNA (Figure 2). However, the continuous PSD of Carp eDNA that we estimated provides simple guidance for overcoming the volume limitations of 0.2 μm filtration by increasing pore size and water volume along an isocline (Figure 3). These isoclines also highlight the potential to exceed the yield of 0.2 μm filtration by using large pore sizes (e.g., >50 μm) and filtering volumes above the isocline. Given the basic processes discussed above regarding the origin, state, and fate of macrobial eDNA, we suspect that these eDNA capture recommendations are applicable beyond Carp. However, as researchers generate eDNA PSDs for additional species and environments, they can easily be compared via Weibull CCDF modeling and our simple equation for calculating isoclines.

The heterogeneous distribution of macrobial eDNA among replicate water samples observed in this study (Figure 2) and others (Jerde *et al.* 2011; Dejean *et al.* 2012; Thomsen *et al.* 2012a; Pilliod *et al.* 2013), suggests an extensive search mode (i.e., large pore size and large water volume) as the default strategy when conducting eDNA-based monitoring of rare species in large habitats. Devices for *in situ* filtration of large volumes, such as inline filters or plankton nets, could maximize water volume and habitat coverage. Our data indicate that capture methods with extremely small size cutoffs (e.g., precipitation, centrifugation, ultrafiltration) are generally unnecessary compared with microfiltration (Figure 2). The >50% increase in non-target eDNA capture (Figure 2) and the severe volume limitations of these methods further reduce their value relative to microfiltration.

Because macrobial eDNA is only a tiny fraction of the eDNA and SPM in water, the risk of interference from non-target eDNA and non-DNA inhibitors increases with sample volume (McDevitt *et al.* 2007; Hata *et al.* 2011; but see Gibson *et al.* 2012). Filtration reduces the risk from non-target eDNA by letting 50% or more pass through (Figure 2), and pre-filtration could reduce the risk from non-DNA inhibitors as SPM was most abundant on 100 or 180 μm filters (Figure 2). In contrast to non-target eDNA, the interference risk from SPM is mitigated by DNA extraction and purification, thus we recommend pre-filtration only if it substantially increases the throughput volume of one’s capture filter. For example, Cary et al. (2006, 2007) applied this ‘band-stop’ filtration approach for eDNA-based surveillance of the invasive diatom *Didymosphenia geminate*.

There is broad interest in research on the reliability of eDNA-based ecological monitoring (Sutherland *et al.* 2013; Zhan *et al.* 2013; Schmidt *et al.* 2013). Particle size analysis, such as that presented here, is one of the many important research avenues needed to better understand the origin, state, and fate of aqueous macrobial eDNA. Our first description of particle size for aqueous macrobial eDNA provides immediate guidance for practitioners and a tested method for researchers. We look forward to future findings that will undoubtedly expand on this initial description by investigating the organismal and environmental determinants of the aqueous eDNA particle size.

## ACKNOWLEDGEMENTS

K.J. Coyne and her lab at the University of Delaware provided generous consultation on qPCR and eDNA extraction methods. R.K. Turner helped make Figure 1. M.A. Renshaw assisted with qPCRs. Laura Arriaga at the Potawatomi Zoo provided sampling access to the carp pond. A.M. Deines provided helpful discussion of PSD modeling. W.L. Chadderton and A.R. Mahon provided helpful discussion of a pilot size fractionation experiment. Funding for this project was provided by the Illinois-Indiana Sea Grant (NA10OAR4170068) and the Great Lakes Restoration Initiative (FY10-S-T0240-O169-2). This work was supported by NSF IGERT grant award #0504495 to the GLOBES graduate training program at the University of Notre Dame. This is a publication of the Notre Dame Environmental Change Initiative.

## Supporting Appendix S1: Estimating model parameters for cumulative PSDs

When modeling the cumulative particle size distribution (PSD) resulting from sequential filtration size fractionation, the explanatory variable (fixed and known) is the pore size of the filter. The response variable is the cumulative proportion of carp eDNA, total eDNA, or SPM retained at that pore size. Particle size fractions ranged from <0.2 μm to >180 μm and the cumulative proportion necessarily ranges from zero to one. Because the final size fraction for each type of aquatic matter (<0.2 μm for Carp eDNA and total eDNA, 0.2–1.0 μm for SPM) necessarily had the cumulative proportion of one, these values were excluded from model estimation.

Because there was no prior data regarding the size distribution of aqueous macrobial eDNA we chose to model it using a complementary cumulative distribution function (CCDF) that is flexible and has been widely used for modeling PSDs of other materials – the Weibull or Rosin/Rammler distribution. We used the CCDF of the Weibull distribution,

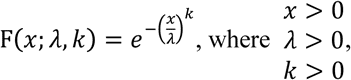

To estimate the model parameter values using maximum likelihood with beta distributed errors (ML-&) we used custom code in Mathematica 8 (Wolfram Research, Champaign, IL, USA). We selected a beta error distribution because the random variable (proportion of total) is continuous and bound between zero and one (Ferrari and Cribari-Neto 2004):

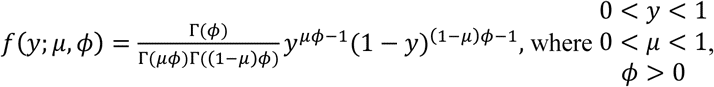

The gamma function is defined as 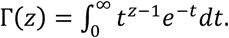

From this model formulation we have a likelihood function of the form,

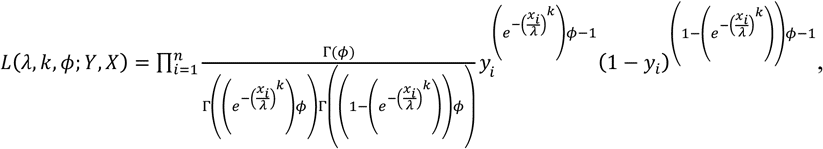

*n* is the number observations. Previous research has used the beta distribution with logistic function for the mean (Polando et al. 2013).

The study had twenty replicate size fractionations (10 pond, 10 lake) and the purpose of the model fitting was to find the best fit parameters under the twenty replicate samples. Thus the likelihood was fit using the multiplication of twenty likelihoods from independent samples (Edwards 1984), such that, *L*(λ, *k*, *ϕ*; *Y*_1_, *X*_1_, *Y*_2_, *X*_2_, …*Y*_20_, *X*_20_) = *L*(λ, *k*, *ϕ*; *Y*_1_, *X*_1_) * *L*(λ, *k*, *ϕ*; *Y*_2_, *X*_2_) * … *L*(λ, *k*, *ϕ*; *Y*_20_, *X*_20_)

All likelihood calculations were conducted in Mathematica 8 (Wolfram Research, Inc. 2010) and maximums were verified graphically.

**Supporting Figure S1.**
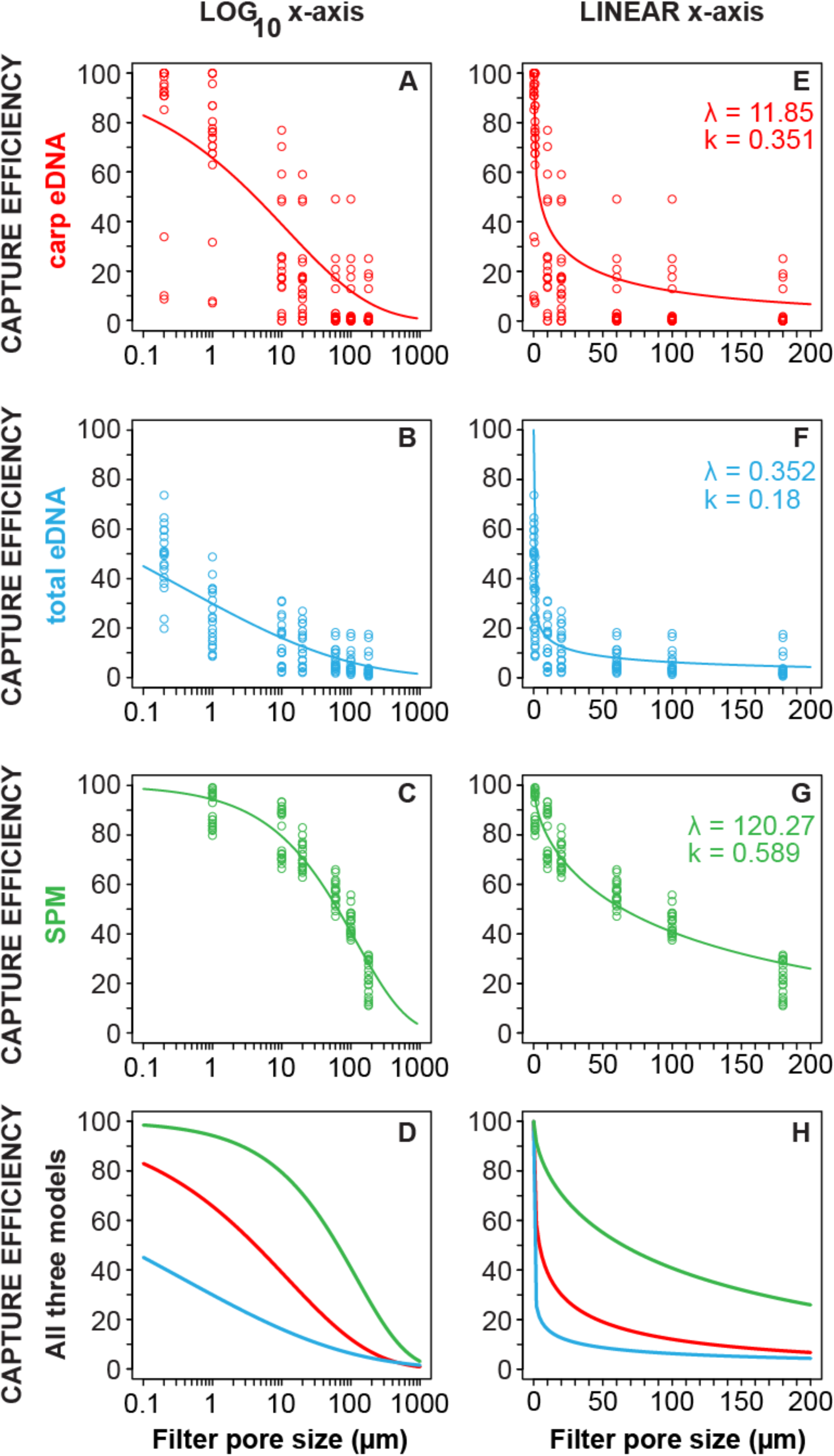
Cumulative size fractionation data and the PSD models (Weibull CCDF) fitted to these data. Pooled lake and pond data are shown for Carp eDNA (A and E), total eDNA (B and F), and SPM (C and G). Panels (D) and (H) show the PSD models for each type of aquatic matter plotted together, without data points. All plots are shown with a log_10_ x-axis (A, B, C, D) and a linear x-axis (E, F, G, H) to allow visualization of both the model fits and the PSD shapes. The estimated scale and shape parameters ($ and k) from the Weibull CCDF model are shown on the plots. Supplemental spring data from the lake were not used for PSD modeling because SPM was not measured in spring.

**Supporting Figure S2.**
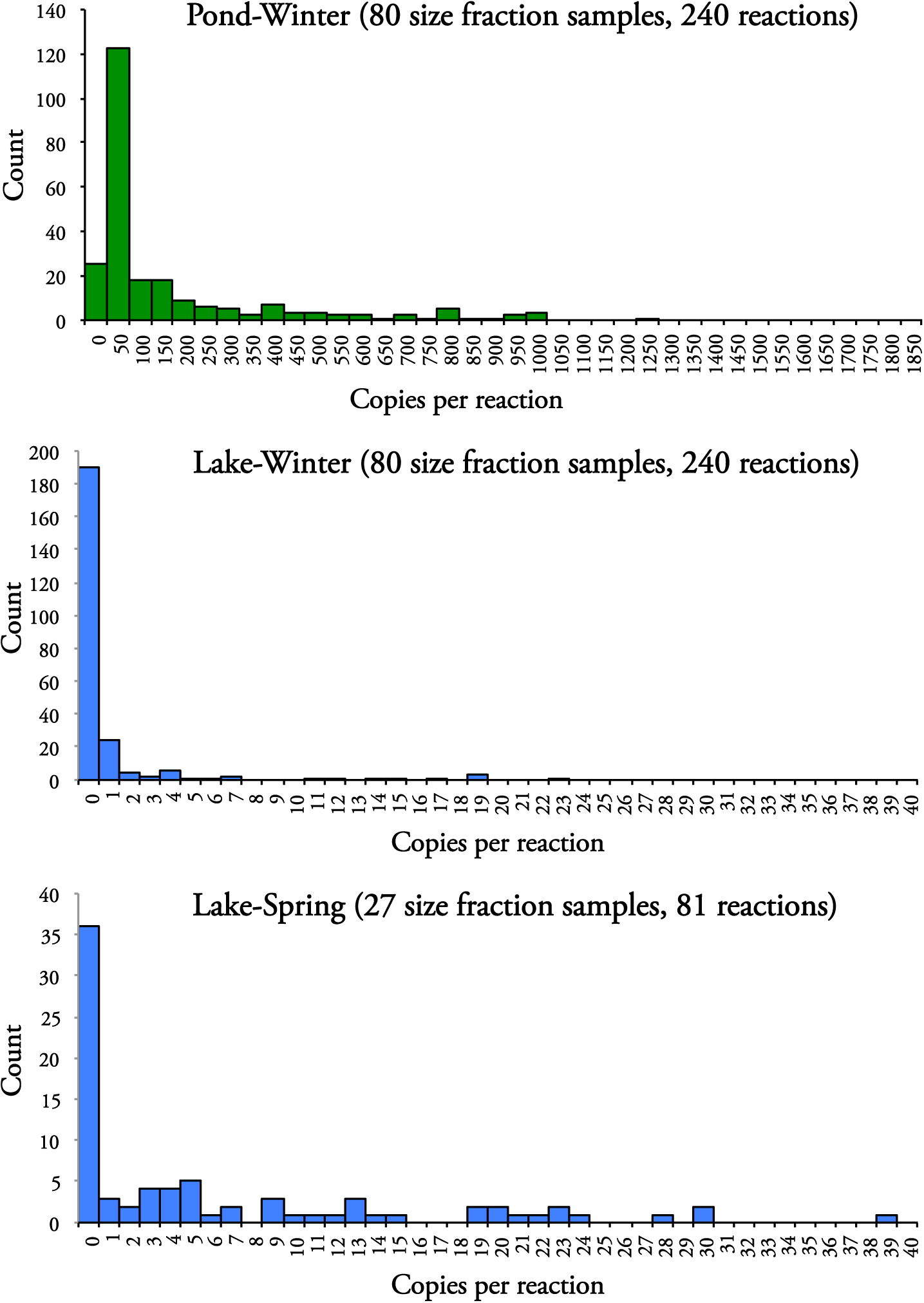
Histograms of qPCR-measured carp eDNA copies per reaction. Note that the x-axis differs between pond and lake histograms.

**Supporting Figure S3.**
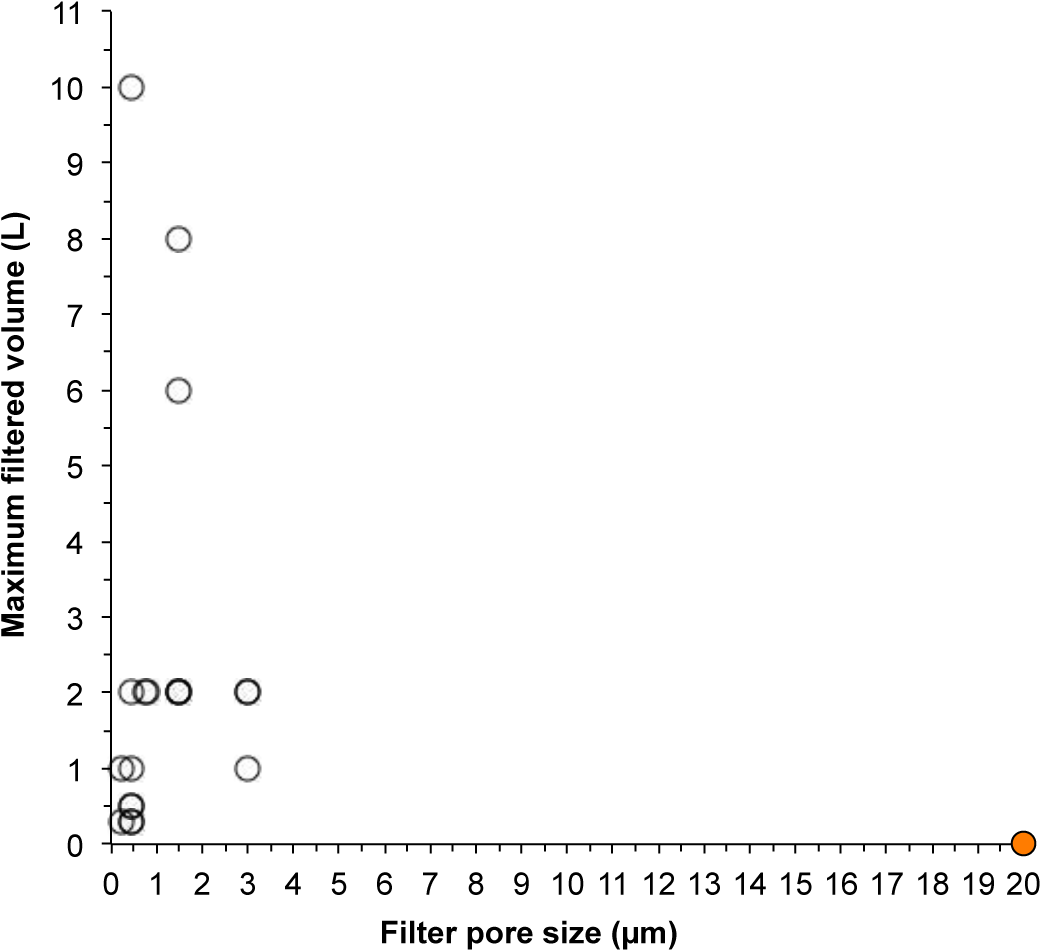
Combinations of filter pore size and water volume used by previous researchers to capture aqueous eDNA from macrofauna. The one filled circle (20 μm pore size) represents a study where the filtered water volume was not reported. Note that filter material and surface area varied across studies. See Table S1 for additional details and published sources.

**Supporting Table S1.**
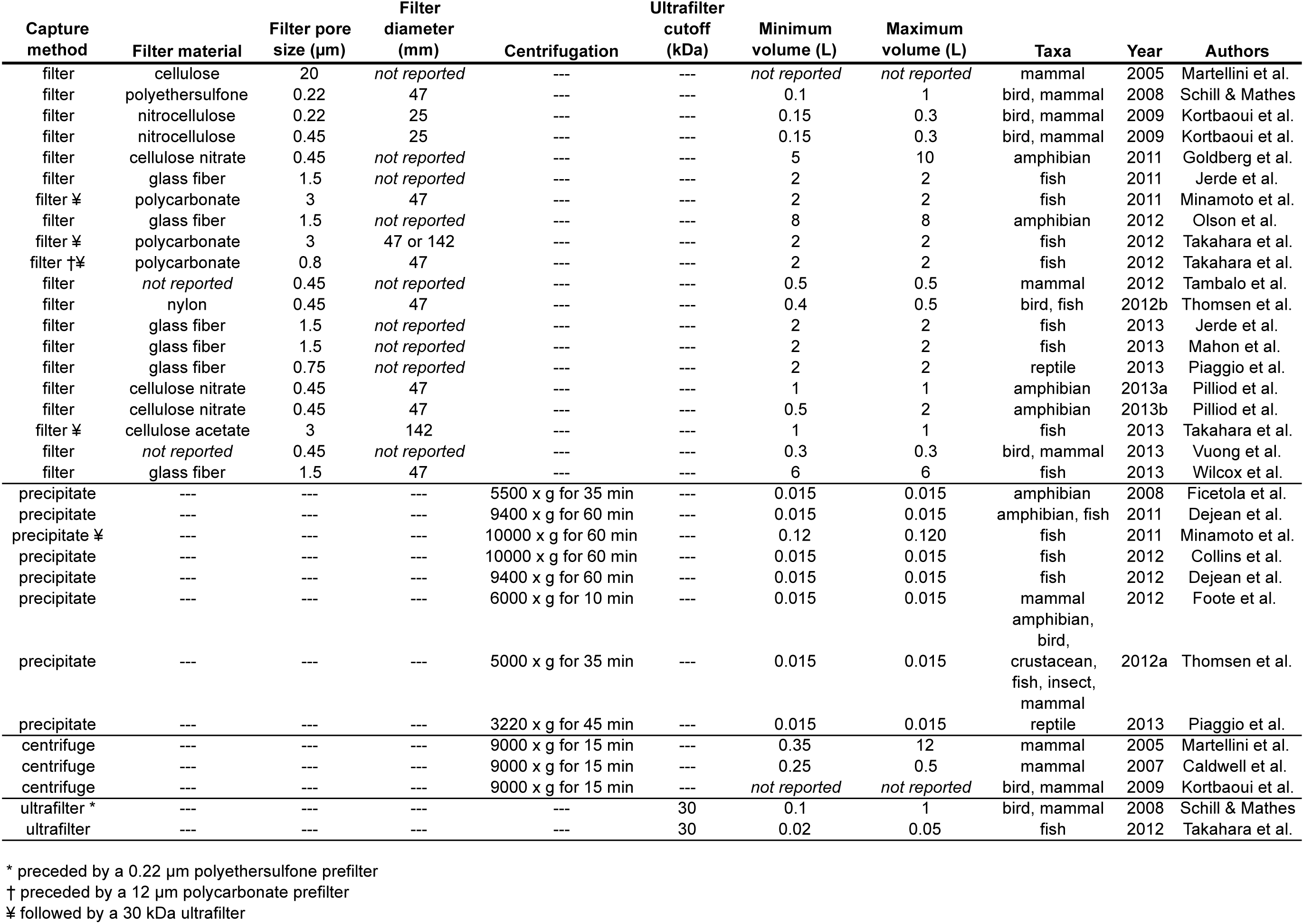
Published methods for capturing aqueous eDNA from macrofauna

## REFERENCES

Bellete, B., Flori, P., Hafid, J., Raberin, H. & Tran Manh Sung, R. (2003). Influence of the quantity of nonspecific DNA and repeated freezing and thawing of samples on the quantification of DNA by the Light Cycler®. Journal of Microbiological Methods, 55, 213–219.

Bloesch, J. (1995). Mechanisms, measurement and importance of sediment resuspension in lakes. Marine and Freshwater Research, 46, 295–304.

Burd, A.B. & Jackson, G.A. (2009). Particle Aggregation. Annual Review of Marine Science, 1, 65–90.

Caldwell, J., Payment, P. & Villemur, R. (2011). Mitochondrial DNA as source tracking markers of fecal contamination. Microbial Source Tracking: Methods, Applications, and Case Studies pp. 229–250 (eds C. Hagedorn, A.R. Blanch & V.J. Harwood), Springer New York, New York, NY.

Cary, S.C., Hicks, B.J., Coyne, K.J., Rueckert, A., Gemmill, C.E.C. & Barnett, C.M.E. (2007). A sensitive genetic-based detection capability for Didymosphenia geminata (Lyngbye) M. Schmidt: Phases Two and Three. Hamilton, New Zealand.

Cary, S.C., Hicks, B.J., Crawford, N.J. & Coyne, K.J. (2006). A sensitive genetic-based detection capability for Didymosphenia geminata. Interim Report. Hamilton, New Zealand.

Collins, R.A., Armstrong, K.F., Holyoake, A.J. & Keeling, S. (2012). Something in the water: biosecurity monitoring of ornamental fish imports using environmental DNA. Biological Invasions.

Coyne, K.J., Handy, S.M., Demir, E., Whereat, E.B., Hutchins, D.A., Portune, K.J., Doblin, M.A. & Cary, S.C. (2005). Improved quantitative real-time PCR assays for enumeration of harmful algal species in field samples using an exogenous DNA reference standard. Limnology and Oceanography-Methods, 3, 381–391.

Dejean, T., Valentini, A., Duparc, A., Pellier-Cuit, S., Pompanon, F., Taberlet, P. & Miaud, C. (2011). Persistence of environmental DNA in freshwater ecosystems. PloS one, 6, e23398.

Dejean, T., Valentini, A., Miquel, C., Taberlet, P., Bellemain, E. & Miaud, C. (2012). Improved detection of an alien invasive species through environmental DNA barcoding: the example of the American bullfrog Lithobates catesbeianus. Journal of Applied Ecology, 49, 953–959.

Douville, M., Gagné, F., Blaise, C. & André, C. (2007). Occurrence and persistence of Bacillus thuringiensis (Bt) and transgenic Bt corn cry1Ab gene from an aquatic environment. Ecotoxicology and environmental safety, 66, 195–203.

Droppo, I.G. (2006). Filtration in Particle Size Analysis. Encyclopedia of Analytical Chemistry (ed R.A. Meyers), pp. 1–16. John Wiley & Sons, Ltd.

Ellison, S.L.R., English, C. a, Burns, M.J. & Keer, J.T. (2006). Routes to improving the reliability of low level DNA analysis using real-time PCR. BMC biotechnology, 6, 33.

Ficetola, G.F., Miaud, C., Pompanon, F. & Taberlet, P. (2008). Species detection using environmental DNA from water samples. Biology Letters, 4, 423–425.

Flindt, R. (2006). Amazing Numbers in Biology, 6th edn. (D. Czeschlik, Ed.). Springer-Verlag Berlin Heidelberg, Heidelberg, Germany.

Foppen, J.W., Orup, C., Adell, R., Poulalion, V. & Uhlenbrook, S. (2011). Using multiple artificial DNA tracers in hydrology. Hydrological Processes, 25, 3101–3106.

Gibson, K.E., Schwab, K.J., Spencer, S.K. & Borchardt, M.A. (2012). Measuring and mitigating inhibition during quantitative real time PCR analysis of viral nucleic acid extracts from large-volume environmental water samples. Water Research, 46, 4281–91.

Goldberg, C.S., Pilliod, D.S., Arkle, R.S. & Waits, L.P. (2011). Molecular detection of vertebrates in stream water: a demonstration using Rocky Mountain tailed frogs and Idaho giant salamanders. PloS one, 6, e22746.

Hata, A., Katayama, H., Kitajima, M., Visvanathan, C., Nol, C. & Furumai, H. (2011). Validation of internal controls for extraction and amplification of nucleic acids from enteric viruses in water samples. Applied and environmental microbiology, 77, 4336–43.

Hedman, J. & Rådström, P. (2013). Overcoming Inhibition in Real-Time Diagnostic PCR. PCR Detection of Microbial Pathogens (ed M. Wilks), pp. 17–48. Humana Press, Totowa, NJ.

Hochmeister, M.N., Budowle, B., Jung, J., Borer, U. V, Comey, C.T. & Dirnhofer, R. (1991). PCR-based typing of DNA extracted from cigarette butts. International Journal of Legal Medicine, 104, 229–233.

Isao, K., Hara, S., Terauchi, K. & Kogure, K. (1990). Role of sub-micrometre particles in the ocean. Nature, 345, 242–244.

Jerde, C.L., Mahon, A.R., Chadderton, W.L. & Lodge, D.M. (2011). “Sight-unseen” detection of rare species using environmental DNA. Conservation Letters, 4, 150–157.

Kibbe, W. a. (2007). OligoCalc: an online oligonucleotide properties calculator. Nucleic acids research, 35, W43–6.

Kortbaoui, R., Locas, A., Imbeau, M., Payment, P. & Villemur, R. (2009). Universal mitochondrial PCR combined with species-specific dot-blot assay as a source-tracking method of human, bovine, chicken, ovine, and porcine in fecal-contaminated surface water. Water Research, 43, 2002–10.

Liu, H. & Chen, Y. (2003). Phylogeny of the East Asian cyprinids inferred from sequences of the mitochondrial DNA control region. Canadian Journal of Zoology, 81, 1938–1946.

Lloyd, K.G., Macgregor, B.J. & Teske, A. (2009). Quantitative PCR methods for RNA and DNA in marine sediments: maximizing yield while overcoming inhibition. FEMS microbiology ecology.

Lodge, D.M., Turner, C.R., Jerde, C.L., Barnes, M. a, Chadderton, L., Egan, S.P., Feder, J.L., Mahon, A.R. & Pfrender, M.E. (2012). Conservation in a cup of water: estimating biodiversity and population abundance from environmental DNA. Molecular Ecology, 21, 2555–8.

Lowe S., Browne M., Boudjelas S., De Poorter M. (2004). 100 of the World’s Worst Invasive Alien Species, a selection from the Global Invasive Species Database. World Conservation Union (IUCN), Auckland, New Zealand. http://www.issg.org/pdf/publications/worst_100/english_100_worst.pdf

Maggi, F. (2013). The settling velocity of mineral, biomineral, and biological particles and aggregates in water. Journal of Geophysical Research: Oceans, 118, 2118–2132.

Martellini, A., Payment, P. & Villemur, R. (2005). Use of eukaryotic mitochondrial DNA to differentiate human, bovine, porcine and ovine sources in fecally contaminated surface water. Water research, 39, 541–8.

Maruyama, F., Tani, K., Kenzaka, T., Yamaguchi, N. & Nasu, M. (2008). Application of Real-Time Long and Short Polymerase Chain Reaction for Sensitive Monitoring of the Fate of Extracellular Plasmid DNA Introduced into River Waters. Microbes and Environments, 23, 229– 236.

Matsui, K., Honjo, M. & Kawabata, Z. (2001). Estimation of the fate of dissolved DNA in thermally stratified lake water from the stability of exogenous plasmid DNA. Aquatic Microbial Ecology, 26, 95–102.

Matsui, K., Ishii, N., Honjo, M. & Kawabata, Z. (2004). Use of the SYBR Green I fluorescent dye and a centrifugal filter device for rapid determination of dissolved DNA concentration in fresh water. Aquatic Microbial Ecology, 36, 99–105.

McDevitt, J.J., Lees, P.S.J., Merz, W.G. & Schwab, K.J. (2007). Inhibition of quantitative PCR analysis of fungal conidia associated with indoor air particulate matter. Aerobiologia, 23, 35–45.

Mifflin, T.E. (2007). Setting up a PCR laboratory. Cold Spring Harbor Protocols pp. 5–14. Cold Spring Harbor Laboratory Press, Cold Spring Harbor, NY, USA.

Minamoto, T., Yamanaka, H., Takahara, T., Honjo, M.N. & Kawabata, Z. (2011). Surveillance of fish species composition using environmental DNA. Limnology, 13, 193–197.

Murgia, M., Pizzo, P., Sandona, D., Zanovello, P., Ruzzuto, R. & Virgilioll, F. Di. (1992). Mitochondrial DNA Is Not Fragmented during Apoptosis. The Journal of Biological Chemistry, 267, 10939–10941.

Ogram, A., Sayler, G.S. & Barkay, T. (1987). The extraction and purification of microbial DNA from sediments. Journal of Microbiological Methods, 7, 57–66.

Piaggio, A.J., Engeman, R.M., Hopken, M.W., Humphrey, J.S., Keacher, K.L., Bruce, W.E. & Avery, M.L. (2013). Detecting an elusive invasive species: A diagnostic PCR to detect Burmese python in Florida waters and an assessment of persistence of environmental DNA. Molecular Ecology Resources.

Pilliod, D., Goldberg, C., Arkle, R.S. & Waits, L.P. (2013). Estimating Occupancy and Abundance of Stream Amphibians Using Environmental DNA from Filtered Water Samples. Canadian Journal of Fisheries and Aquatic Sciences, 1130, 1123–1130.

Poté, J., Mavingui, P., Navarro, E., Rosselli, W., Wildi, W., Simonet, P. & Vogel, T.M. (2009). Extracellular plant DNA in Geneva groundwater and traditional artesian drinking water fountains. Chemosphere, 75, 498–504.

Reynolds, R.A., Stramski, D., Wright, V.M. & Woźniak, S.B. (2010). Measurements and characterization of particle size distributions in coastal waters. Journal of Geophysical Research: Oceans, 115, C08024.

Rickwood, D. & Chambers, J.A. (1981). Evidence for protected regions of DNA in the mitochondrial nucleoid of Saccharomyces cerevisiae. Fems Microbiology Letters, 12, 187–190.

Robison, B.H. & Bailey, T. (1981). Sinking rates and dissolution of midwater fish fecal matter. Marine Biology, 65, 135–142.

Rosin, P. & Rammler, E. (1933). The Laws Governing the Fineness of Powdered Coal. Journal of the Institute of Fuel, 7, 29–36.

Sabir, I.H., Torgersen, J., Haldorsen, S. & Aleström, P. (2002). DNA tracers with information capacity and high detection sensitivity tested in groundwater studies. Hydrogeology Journal, 7, 264–272.

Saeki, K., Ihyo, Y., Sakai, M. & Kunito, T. (2011). Strong adsorption of DNA molecules on humic acids. Environmental Chemistry Letters, 9, 505–509.

Schmidt, B.R., Kéry, M., Ursenbacher, S., Hyman, O.J. & Collins, J.P. (2013). Site occupancy models in the analysis of environmental DNA presence/absence surveys: a case study of an emerging amphibian pathogen. Methods in Ecology and Evolution, 4, 646–653.

Schrader, C., Schielke, a, Ellerbroek, L. & Johne, R. (2012). PCR inhibitors - occurrence, properties and removal. Journal of Applied Microbiology, 113, 1014–26.

Sheldon, R.W. (1972). Size separation of marine seston by membrane and glass-fiber filters. Limnology and Oceanography, 17, 494–498.

Sutherland, W.J., Bardsley, S., Clout, M., Depledge, M.H., Dicks, L.V, Fellman, L., Fleishman, E., Gibbons, D.W., Keim, B., Lickorish, F., Margerison, C., Monk, K.A, Norris, K., Peck, L.S., Prior, S.V., Scharlemann, J.P.W., Spalding, M.D. & Watkinson, A.R. (2013). A horizon scan of global conservation issues for 2013. Trends in Ecology & Evolution, 28, 16–22.

Suzuki, H., Daimon, M., Awano, T., Umekage, S., Tanaka, T. & Kikuchi, Y. (2009). Characterization of extracellular DNA production and flocculation of the marine photosynthetic bacterium Rhodovulum sulfidophilum. Applied Microbiology and Biotechnology, 84, 349–56.

Takahara, T., Minamoto, T. & Doi, H. (2013). Using environmental DNA to estimate the distribution of an invasive fish species in ponds. PloS one, 8, e56584.

Takahara, T., Minamoto, T., Yamanaka, H., Doi, H. & Kawabata, Z. (2012). Estimation of fish biomass using environmental DNA. PloS ONE, 7, e35868.

Tepper, C.G. & Studzinski, G.P. (1993). Resistance of mitochondrial DNA to degradation characterizes the apoptotic but not the necrotic mode of human leukemia cell death. Journal of Cellular Biochemistry, 52, 352–61.

Thompson, D.E., Rajal, V.B., De Batz, S. & Wuertz, S. (2006). Detection of Salmonella spp. in water using magnetic capture hybridization combined with PCR or real-time PCR. Journal of Water and Health, 4, 67–75.

Thomsen, P.F., Kielgast, J., Iversen, L.L., Wiuf, C., Rasmussen, M., Gilbert, M.T.P., Orlando, L. & Willerslev, E. (2012a). Monitoring endangered freshwater biodiversity using environmental DNA. Molecular Ecology, 21, 2565–2573.

Thomsen, P.F., Kielgast, J., Iversen, L.L., Møller, P.R., Rasmussen, M. & Willerslev, E. (2012b). Detection of a diverse marine fish fauna using environmental DNA from seawater samples. PloS one, 7, e41732.

Widmer, F., Seidler, R.J., Donegan, K.K. & Reed, G.L. (1997). Quantification of transgenic plant marker gene persistence in the field. Molecular Ecology, 6, 1–7.

Wilcox, T.M., McKelvey, K.S., Young, M.K., Jane, S.F., Lowe, W.H., Whiteley, A.R. & Schwartz, M.K. (2013). Robust detection of rare species using environmental DNA: the importance of primer specificity. PloS one, 8, e59520.

Willerslev, E., Hansen, A.J., Binladen, J., Brand, T.B., Gilbert, M.T.P., Shapiro, B., Bunce, M., Wiuf, C., Gilichinsky, D.A. & Cooper, A. (2003). Diverse plant and animal genetic records from Holocene and Pleistocene sediments. Science, 300, 791–795.

Wotton, R.S. & Malmqvist, B. (2001). Feces in Aquatic Ecosystems. Bioscience, 51, 537–544.

Zhan, A., Hulák, M., Sylvester, F., Huang, X., Adebayo, A.A., Abbott, C.L., Adamowicz, S.J., Heath, D.D., Cristescu, M.E. & MacIsaac, H.J. (2013). High sensitvity of 454 pyrosequencing for detection of rare species in aquatic communities. Methods in Ecology and Evolution, 4, 558– 565.

Zobeck, T.M., Gill, T.E. & Popham, T.W. (1999). A two-parameter Weibull function to describe airborne dust particle size distributions. Earth Surface Processes and Landforms, 24, 943–955.

## REFERENCES for Supporting Appendix S1

Edwards, Anthony WF. Likelihood. CUP Archive, 1984.

Ferrari, Silvia, and Francisco Cribari-Neto. “Beta regression for modelling rates and proportions.” Journal of Applied Statistics 31, no. 7 (2004): 799–815.

Polando, Rachel, Upasna Gaur Dixit, Cristina R. Carter, Blake Jones, James P. Whitcomb, Wibke Ballhorn, Melissa Harintho, Christopher L. Jerde, Mary E. Wilson, and Mary Ann McDowell. “The roles of complement receptor 3 and Fc’ receptors during Leishmania phagosome maturation.” Journal of leukocyte biology (2013)

